# Correlation between Serum Kisspeptin and Spermatogenic Function in Men

**DOI:** 10.1101/810572

**Authors:** Hongling Yu, Jin Liu, Yilong Han, Chao Chen, Fanwei Meng

**Affiliations:** Infertility Center of Qilu Hospital of Shandong University, Jinan, Shandong,China

**Author notes:** Corresponding author: Fanwei Meng. (FW M). These authors contributed equally to this work. Author contributions: Conceived and designed the experiments: Meng Fanwei, Performed the experiments: Yu Hongling, Analyzed the data: Meng Fanwei; Yu Hongling, Contributed reagents/materials/analysis tools: Han Yilong; Liu Jin; Chenchao.

**Keywords:** FSH, kisspeptin, LH, male infertility, testosterone

## Abstract

Kisspeptin along with its receptor GPR54 or KISS1R regulates the secretion of hormones involved in the hypothalamic-pituitary-testicular axis, which is one of the contributing factors of adolescent spermatogenesis. This study aimed to investigate the correlation between serum kisspeptin concentration and spermatogenic function, and its predictive value in azoospermia. We retrospectively analyzed data of 196 males who visited the Reproductive Medicine Center of Qilu Hospital of Shandong University from June–November 2018; 20 were fertile and 176 were infertile. The following semen tests were performed: serum kisspeptin level by enzyme immunoassay; and levels of follicle-stimulating and luteinizing hormones, and testosterone by chemiluminescence assay. Percutaneous testicular sperm aspiration was performed on males with azoospermia. Subjects were divided into 5 groups: azoospermia (group A: 22 men with obstructive azoospermia; group B: 54 men with non-obstructive azoospermia), oligospermia (group C: 56 men), infertility with normal semen concentration (group D: 44 men), and fertility with normal semen concentration (group E: 20 men). Kisspeptin levels in the fertile group were higher than those in the infertile group. Levels of serum hormones, testosterone, and kisspeptin correlated with sperm concentration, with the strongest correlation between kisspeptin and sperm concentration (correlation coefficient=0.692). Levels of kisspeptin in obstructive and non-obstructive azoospermia groups were analyzed using Receiver-Operating-Curve analysis. A serum kisspeptin level ≥80.655 was classified as obstructive azoospermia; otherwise, the classification was non-obstructive azoospermia. Serum kisspeptin levels in the fertility group were significantly higher than that in the infertility group; this suggests kisspeptin may be associated with male fertility. Moreover, kisspeptin had a stronger correlation with sperm concentration than the hormones. A serum kisspeptin level of 80.655 can be used to differentiate obstructive and non-obstructive azoospermia.

## Introduction

Infertility is a social problem in modern society, and the incidence in the reproductive population is 10-20%. Half of all cases are attributed directly or indirectly to male infertility ^[1]^. Oligospermia, asthenospermia, and azoospermia are the main causes of male infertility; approximately 10-20% of males with infertility have azoospermia ^[2]^ and 60% have non-obstructive azoospermia. Azoospermia is caused by testicular spermatogenic dysfunction that develops for various reasons, and it is difficult to find any sperm in males with azoospermia by conventional methods.

The functional status of testicular tissues is the main factor that affects sperm quality. The generation and apoptosis of germ cells are precisely regulated by endocrine and paracrine hormones ^[3]^. In spermatogenesis and sperm development, the hypothalamic-pituitary-gonadal (HPG) axis regulates the secretion of male sex hormones, which affects the generation and maturation of sperm through different pathways; there are associations of positive and negative feedback among these hormones ^[4, 5]^. In 1996, Lee et al^[6]^. discovered and named the *KISS-1* gene while studying the different metastatic activity of human melanoma cells using subtractive hybridization. The *KISS-1*gene is located on chromosome 1q32-41 in humans and contains 4 exons and 3 introns. The first two exons are not translated, and the 5’ end of the third exon contains 38 non-coding base pairs followed by a transcription start site and 100 coding base pairs. The fourth exon contains 332 coding base pairs and a stop codon ^[7]^. The *KISS-1* gene encodes Metastin (tumor metastasis suppressor), and its protein product is called kisspeptin ^[8]^. The expression level of the *KISS-1* gene is highest in the human placenta, followed by the testis, pancreas, small intestine, and liver ^[9]^. In the brain, the mRNA of *KISS-1* is discretely distributed in the central nervous system, including the basilar part of the nerve center and hypothalamus ^[10]^. The mRNA encodes a protein that contains 145 amino acid residues—kisspeptin, which plays a biological role. Kisspeptin regulates the secretion of gonadotropin releasing hormone (GnRH) in the hypothalamus and pituitary gonadotropin through its G-protein coupled receptor 54, which participates in the regulation of mammalian sexual maturity and development of the reproductive system during adolescence ^[8, 11]^.

In-depth clinical research and analysis of the proper evaluation of spermatogenic function in infertile males with existing examination methods, appropriate treatment measures for next steps, and avoidance of unnecessary testicular biopsy would be beneficial. This study retrospectively analyzed the sperm parameters and concentrations of serum sex hormones and kisspeptin in 176 infertile and 20 fertile men. We aimed to elucidate the correlation between serum kisspeptin level and male spermatogenic function, and its predictive value in men with azoospermia.

## Materials and methods

### Research subjects

The enrolled subjects were 196 males who visited the Infertility Center of Qilu Hospital of Shandong University from June to November 2018. The age of the subjects ranged from 22 to 50 years. After a physical examination and collection of medical history, the following routine semen tests were performed according to the WHO laboratory manual for the examination and processing of human semen (2010): Serum kisspeptin level by enzyme immunoassay; and levels of follicle-stimulating hormone (FSH), luteinizing hormone (LH), and testosterone (T) by chemiluminescence assay. Percutaneous testicular sperm aspiration was performed on males with azoospermia. Subjects were divided into 5 groups according to their sperm concentration: the azoospermia group (group A: 22 men with obstructive azoospermia; group B: 54 men with non-obstructive azoospermia), the oligospermia group (group C: 56 men), the infertility group with normal semen concentration (group D: 44 men), and the fertility group with normal semen concentration (group E: 20 men). The correlation between both serum sex-hormone levels and kisspeptin, and sperm concentration was analyzed, and the predictive value of kisspeptin for spermatogenic function in men with azoospermia was studied.

All subjects had been physically healthy and diseases, including tuberculosis, asthma, liver and kidney disease, hypertension, and diabetes, were excluded. The inclusion criteria were as follows: (1) infertile males aged 22–50 years who had regular sexual activity; (2) males with normal sexual function who could collect their semen by masturbation; and (3) males who complied with the laboratory sampling standards. The exclusion criteria were as follows: (1) males with a chromosomal abnormality; (2) males who were unable to collect their semen by masturbation; (3) males who were mentally overstressed; (4) males who were exposed to toxic substances that could affect fertility, such as pesticide-factory workers, welders, painters; (5) males with conditions that could affect fertility, such as hypospadias, severe varicocele, cryptorchidism, tumors, male sterilization, and testicular torsion; and (6) males with severe inflammation of their accessory sex glands.

This study was approved by the Ethics Committee of Qilu Hospital of Shandong University, and all subjects signed an informed consent.

### Research methods

1. Semen examination: Semen samples were collected and processed according to the WHO laboratory manual for the examination and processing of human semen (2010) ^[12]^. The total sperm count, concentration, percentage of forward progression, and percentage of sperm motility were determined by the computer-assisted semen analysis operating system using a Makler counting cell that was loaded with the samples. For each semen sample with a sperm concentration <1×10^6^/mL, the number of sperm in 20 different high-power microscopic fields were counted to calculate the concentration of sperm.
2. Determination of the levels of serum kisspeptin and sex hormones: The upper serum was taken from coagulated blood samples for testing. The Cobas e 601 (Roche, Germany) electrochemiluminescence device was used to determine the concentrations of serum FSH, LH, estradiol (E2), prolactin (PRL), and T. The electrochemiluminescence microparticle immunoassay kit was provided by Roche, Germany. The serum kisspeptin concentration was measured by an enzyme-linked immunosorbent assay.
3. Testicular sperm aspiration: Local anesthesia with 0.5% lidocaine was administered. Percutaneous puncture of the testes was then performed with a specialized puncture needle. The tissue obtained was pulverized and observed under a microscope for sperm; in addition, some testicular tissues were sent for a pathological examination.

### Statistical analysis

All statistical analyses were performed using SPSS 16.0 software. Continuous data were expressed as the mean ± standard error, the differences among groups were compared by one-way analysis of variance (ANOVA), and the pairwise comparison between groups was performed by the Tukey method. The correlation between each parameter and spermatogenic function was expressed by the Pearson correlation coefficient. The cut-off value of the level of kisspeptin for predicting the spermatogenic function of males with azoospermia was determined by the Receiver Operating Characteristic (ROC) method. The stepwise regression method was used to establish a regression model for the prediction of spermatogenic function.

## Results

### Levels of serum E2, FSH, LH, T, kisspeptin, and age in the different groups

Table 1 shows the levels of serum E2, FSH, LH, T, kisspeptin, and age in the different groups. The univariate ANOVA showed statistical differences of serum E2, FSH, LH, T, and kisspeptin levels among the five groups.

**Table 1.** Levels of serum E2, FSH, LH, T, kisspeptin, and age in the different groups

| Parameters | A<br>(n=22) | B<br>(n=54) | C<br>(n=56) | D<br>(n=44) | E<br>(n=20) | F | P-value |
| --- | --- | --- | --- | --- | --- | --- | --- |
| Age | 28.73±2.07 | 29.11±0.96 | 29.46±1.2 | 29.41±1.04 | 32.8±2.1 | 0.890 | 0.473 |
| E2 | 29.36±3.71 | 27.41±3.23 | 32.85±2.57 | 29.16±2.66 | 10.42±0.82 | 5.153 | 0.001 |
| FSH | 6.53±2.2 | 14.45±2.84 | 8.01±0.98 | 3.79±0.2 | 4.3±0.41 | 5.576 | <0.001 |
| LH | 4.5±0.82 | 10.15±1.99 | 6.84±0.58 | 3.67±0.22 | 3.69±0.3 | 4.920 | 0.001 |
| PRL | 10.92±0.6 | 15.42±2.4 | 18.6±5.93 | 12.47±0.94 | 12.01±0.35 | 0582 | 0.676 |
| T | 3.35±0.55 | 2.82±0.48 | 4.68±0.28 | 4.26±0.28 | 3.84±0.14 | 4.292 | 0.003 |
| KISS | 150.82±16.64 | 46.33±3.79 | 73.45±6.96 | 100.07±10.71 | 196.29±15.07 | 33.051 | <0.001 |

It shows a significant decrease in the E2 concentration (ng/mL) in the non-obstructive azoospermia group (27.41 ± 3.23, P < 0.001). The serum E2 levels in the obstructive azoospermia group, the oligospermia group, the infertility group with normal semen concentration, and the fertility group with normal semen concentration were 29.36 ± 3.71, 32.85 ± 2.57, 29.16 ± 2.66, and 10.42 ± 0.82, respectively.

It also shows a significant increase in the FSH concentration (ng/mL) in the non-obstructive azoospermia group (14.45 ± 2.84, P < 0.001). The serum FSH levels in the obstructive azoospermia group, the oligospermia group, the infertility group with normal semen concentration, and the fertility group with normal semen concentration were 6.53 ± 2.2, 8.01 ± 0.98, 3.79 ± 0.2, and 4.3 ± 0.41, respectively.

Table 1 shows a significant increase in LH concentration (ng/mL) in the non-obstructive azoospermia group (10.15 ± 1.99, P < 0.001). The serum LH levels in the obstructive azoospermia group, the oligospermia group, the infertility group with normal semen concentration, and the fertility group with normal semen concentration were 4.5 ± 0.82, 6.84 ± 0.58, 3.67 ± 0.22, and 3.69 ± 0.3, respectively.

It shows a significant decrease in the T concentration (ng/mL) in the non-obstructive azoospermia group (2.82 ± 0.48, P < 0.05). The serum T levels in the obstructive azoospermia group, the oligospermia group, the infertility group with normal semen concentration, and the fertility group with normal semen concentration were 3.35 ± 0.55, 4.68 ± 0.28, 4.26 ± 0.28, and 3.84 ± 0.14, respectively.

It also shows a significant decrease in the kisspeptin concentration (ng/mL) in the non-obstructive azoospermia group (46.33 ± 3.79, P < 0.001). The serum kisspeptin levels in the obstructive azoospermia group, the oligospermia group, the infertility group with normal semen concentration, and the fertility group with normal semen concentration were 150.82 ± 16.64, 73.45 ± 6.96, 100.07 ± 10.71, and 196.29 ± 15.07, respectively.

### Correlations between blood parameters and spermatogenic function

The Pearson correlation coefficients show that E2, FSH, LH, and kisspeptin are statistically correlated to spermatogenic function, and the correlation between kisspeptin and spermatogenic function (correlation coefficient = 0.692) is the strongest (Table 2).

**Table 2.** Correlations between blood parameters and spermatogenic function

|  | E2 | FSH | LH | T | kisspeptin |
| --- | --- | --- | --- | --- | --- |
| Pearson correlation coefficient | -0.318 | -0.291 | -0.286 | 0.040 | 0.692 |
| P-value | <0.001 | 0.004 | 0.004 | 0.695 | <0.001 |

The Pearson correlation coefficients show that E2, FSH, LH, and kisspeptin are statistically correlated to spermatogenic function, and the correlation between kisspeptin and spermatogenic function (correlation coefficient = 0.692) is the strongest (Table 2).

### Predictive value of kisspeptin for the spermatogenic function of males with azoospermia (obstructive vs. non-obstructive)

ROC-curve analysis was performed. The sensitivity, specificity, and Youden index of the different cut-off values, and the area under the curve (AUC) of the ROC curves were calculated. A final cut-off value was designated when the Youden index was at maximum (Table 3).

**Table 3.** Diagnosis of obstructive vs. non-obstructive azoospermia by kisspeptin

| Category | Cut-off value | AUC | Sensitivity | Specificity | Youden index |
| --- | --- | --- | --- | --- | --- |
| obstructive vs.<br>non-obstructive | 80.655 | 0.997 | 1 | 0.963 | 0.963 |

ROC-curve analysis was performed. The sensitivity, specificity, and Youden index of the different cut-off values, and the area under the curve (AUC) of the ROC curves were calculated. A final cut-off value was designated when the Youden index was at maximum (Table 3).

Kisspeptin has a predictive value for the spermatogenic function of males with azoospermia. When a male’s kisspeptin value is ≥80.655, their condition is classified as obstructive azoospermia; otherwise, it is classified as non-obstructive azoospermia.

### Regression model

Age and the levels of E2, FSH, LH, T, and kisspeptin were included in the regression model for predicting spermatogenic function. The results are shown in Table 4.

**Table 4.** Regression prediction model

| Variables | $\beta$ | t | P-value |
| --- | --- | --- | --- |
| Constant term | 15.454 | 2.094 | 0.039 |
| Kisspeptin | 0.189 | 4.798 | <0.001 |
| E2 | -0.396 | -2.460 | 0.016 |
| FSH | -0.556 | -2.242 | 0.027 |

The final equation for the model is Y=15.454+0.189×kisspeptin-0.396×E2-0.556×FSH

## Discussion

Spermatogenic ability is one of the important factors that affects semen quality. Previous studies have shown that sperm generation and development are primarily regulated by the HPG axis. The hypothalamus’ pulsatile secretion of gonadotropin-releasing hormone (GnRH) acts on the pituitary gland to promote synthesis and secretion of FSH. FSH then acts on the seminiferous tubules and Sertoli cells to promote the development of spermatocytes of all stages and sperm maturation ^[13, 14]^.

Previous studies have shown that males with precocious puberty have higher levels of serum kisspeptin compared to those of a control group of the same age ^[15–17]^. Recent studies have also reported that the plasma levels of kisspeptin, LH, and FSH in fertile men are significantly higher than those in men with hypogonadism ^[18]^. Moreover, studies have shown that an injection of kisspeptin-54 powerfully stimulates the release of LH, FSH, and T ^[19]^. Therefore, kisspeptin is clearly involved in regulating the hormone secretion of the HPG axis and plays an important role in sexual maturity and development of the reproductive system in mammals. Muhammad ^[20]^ found that the level of serum kisspeptin in infertile men was significantly lower than that in a fertile control group; we reached the same conclusion in this study.

Madani et al. ^[21]^ believed that a combined determination of FSH level and testicular size could replace testicular biopsy to predict the presence or absence of spermatogenesis in males with azoospermia. Studies have shown that serum inhibin-B (INH-B) is associated with sperm count, testicular volume, and testicular pathological scores, and it can be used as a marker of testicular spermatogenic function ^[22, 23]^. Most studies have used serum INH-B levels to predict testicular biopsy outcomes ^[24, 25]^, but some studies have questioned the reliability of serum INH-B levels to predict spermatogenic function ^[26, 27]^. Mitchell et al. ^[28]^ believed that even seminal plasma INH-B could not predict the outcome of testicular biopsies. For serum anti-mullerian hormone (AMH), there is currently no consensus as to whether it can be used as an indicator of spermatogenesis ^[29–31]^. Shanthi et al. ^[23]^ reported that serum AMH concentration in males with non-obstructive azoospermia was significantly lower than that in males with obstructive azoospermia and normal sperm concentrations. Al-Qahtani et al. ^[32]^ and Appasamy et al. ^[33]^ reported that serum AMH concentration in infertile males was significantly lower than that of normal males. However, some studies have reported no statistically significant differences in serum AMH levels between males with oligospermia and those with normal sperm concentrations, or between infertile men and fertile men ^[34, 35]^. We explored the correlation between serum levels of sex hormones and kisspeptin, and sperm concentration in infertile and fertile men to determine if the serum kisspeptin level could be used as an indicator of male spermatogenesis. We discovered that E2, FSH, LH, T, and kisspeptin are all correlated to sperm concentration. In males with non-obstructive azoospermia, E2 and T were significantly decreased, and FSH and LH were significantly increased. The level of kisspeptin increased as the sperm concentration increased. After calculating the Pearson correlation coefficient, E2, FSH, LH, and kisspeptin were statistically correlated to sperm concentration, and the correlation between kisspeptin and sperm concentration (correlation coefficient = 0.692) was stronger than the correlation between FSH or LH and sperm concentration. We conclude that the level of kisspeptin can predict male spermatogenesis.

We also studied the serum levels of kisspeptin in males with obstructive and non-obstructive azoospermia. ROC-curve analysis was performed and a cut-off value was designated at the maximum of the Youden index. The level of kisspeptin was of diagnostic value for obstructive azoospermia. When the AUC is 0.997, the optimal cut-off value is 80.655. In other words, when a male’s serum kisspeptin level is ≥80.655, the condition is classified as obstructive azoospermia; otherwise, it is classified as non-obstructive azoospermia.

The application of microsurgery for collecting sperm has enabled more males with non-obstructive azoospermia to collect their sperm through surgery and fulfill their fertility needs with the assistance of in-vitro fertilization technology. We are currently collecting more samples of serum and testicular tissue from infertile males, especially those with azoospermia, to conduct research on kisspeptin. We look forward to providing better tools to identify obstructive and non-obstructive azoospermia and to predict the fertility and infertility of men in the future.

## Author contributions

Conceived and designed the experiments: Meng Fanwei

Performed the experiments: Yu Hongling

Analyzed the data: Meng Fanwei; Yu Hongling

Contributed reagents/materials/analysis tools: Han Yilong; Liu Jin; Chenchao

## Acknowledgement

Acknowledgement our colleagues of QILU hospital worked for this article.

